# Winter wheat adapts to environmental pH by changing H^+^ net flux in roots at the seedling stage

**DOI:** 10.1101/2020.10.27.356840

**Authors:** Guangtao Wang, Suwei Feng, Weihua Ding, Tiezhu Hu, Zhengang Ru

## Abstract

Changes in rhizosphere pH play an important role in wheat growth. To investigate the relationship between changes in rhizosphere pH and the growth of winter wheat roots and to explore the regulatory mechanism of acid and alkali resistance in winter wheat roots, the semi-winter wheat varieties Aikang 58 (AK58) and Bainong 4199 (BN4199) were used as materials for hydroponic experiments. Three pH levels (4.0, 6.5, and 9.0, with 6.5 as control) were applied during the wheat seedling stage. The results showed that the shoot and root biomass of the plants significantly decreased compared with the control under acid-base stress, with a more significant decrease with acid stress than alkali stress. Compared with the control, the root/shoot ratio increased under alkali stress and decreased under acid stress. The wheat root system showed H^+^ net efflux at pH 6.5 and 9.0, and the H^+^ net efflux rate at pH 4.0 was significantly lower than the control. The root activity of wheat was higher than the control at pH 9.0 and lower at pH4.0. The change of root pH was showed pH 4.0 < pH 6.5 < pH 9.0. Correlation analysis showed that changes in H^+^ net flux were significantly positively correlated to root activity and root pH. The H^+^ efflux rate and root activity of BN4199 were higher than AK58 under acid and alkali stress, and the root/shoot ratio was relatively high, indicating strong acid and alkali resistance. We conclude that wheat could adapt to poor acid-base environments by adjusting root H^+^ net flux, and in practice, the root/shoot ratio could be used as index for the rapid determination of acid-base tolerance in wheat at the seedling stage.

## Introduction

Wheat (*Triticum aestivum* L.) is an important cereal crop globally, and its growth and yield formation are largely affected by soil environmental conditions [1,2], particularly the recent intensification of soil acidification, which has severely affected food production and sustainable agricultural development [3,4]. The optimal soil pH for wheat growth is 6.5–7.0, and excessively acidic or alkaline soil may restrict the growth and development of wheat. Soil pH can affect the absorption of different ions by crops [5], thus affecting the overall growth status of wheat. Although reasonable water and soil management and chemical regulation play important roles in improving soil chemical properties [6], large-scale application improvements are relatively time-consuming and costly [7,8]. Therefore, cultivating acid- and alkali-tolerant wheat varieties is the most economical and effective method in addressing wheat propagation in acidic or alkaline soil.

Soil pH is largely influenced by the H^+^ concentration in the soil solution. In wheat production, the root system is in direct contact with the soil environment and is an important organ for the absorption and transport water and nutrients [9]. Hinsinger et al. [10] suggested that root-induced proton efflux or influx could respectively result in a significant decrease or increase in rhizosphere pH, usually up to ± 0.1 to 1 pH unit, and in some cases as high as ± 2–3 pH unit. Cu et al. [11] showed that wheat monocultures raised soil pH by 0.8 unit, while lupin monoculture lowered it by 0.3 pH unit, and mixed cultures result in soil pH that is intermediate of the two monocultures. In a strongly acidic, Cu-contaminated soil, the rhizosphere pH of wheat increased over 6 mm in soil, reaching up to +2.8 pH units close to the root mat surface [11]. Root-induced changes in rhizosphere pH will change the availability of nutrients in soil [12], thereby affecting plant growth and development. At the rhizosphere range, it has been demonstrated that protons are released by roots to compensate for the excessive absorption of cations by plants over anions [13]. The concentration of protons determines the pH of the solution, and H^+^-ATPase activity helps maintain the pH of cells and provides an H^+^ electrochemical gradient for the influx or efflux of ions into or from the cells [14]. Xue et al. [15] suggested that enhancing the vacuolar Na^+^/H^+^ antiporter protein levels could improve salt tolerance in wheat in saline soils. Bouthour et al. [16] suggested that the NADPH circulatory system is associated with enhanced salt tolerance in wheat. In addition, soil acidification may cause soil aluminum (Al) toxicity and heavy metal pollution, which will cause active Al toxicity in wheat [17] as well as Cu [18], Fe [19], Mn [20], Pb [21], Cr [22], and other heavy metal poisoning. In addition, soil salinization can cause damage to wheat such as water stress, ion toxicity, and oxidative stress, thus affecting growth [23]. Borzouei et al. [24] found that wheat varieties with strong tolerance have better growth characteristics, and the yield of tolerant wheat varieties is higher than that of sensitive varieties [25]. Therefore, it is of great significance to select acid-alkali-resistant wheat varieties for stress tolerance breeding.

Wherrett et al. [26] found that the rhizosphere environment of wheat would affect changes in root H^+^ net flux. However, studies on changes in H^+^ net flux in wheat roots under different pH conditions are limited. Because roots are the first plant parts that respond to stress, it is highly relevant to study the effect of different pH levels on wheat growth using roots, including its underlying mechanism of resistance to acid and alkali stress. Therefore, hydroponics was used to simulate different rhizosphere pH environments, analyze biomass, root/shoot ratio, root H^+^ net flux, root activity, and root pH of winter wheat and to explore the root regulatory mechanism of wheat to adapt to different rhizosphere pH levels to provide a reference for identifying acid- and alkali-resistant varieties and yield improvement.

## Materials and methods

### Plant materials and treatment

Two winter wheat (*Triticum aestivum* L.) varieties, Aikang58 (AK58) and Bainong4199 (BN4199), which are widely cultivated in the North China Plain, were used in this study. Wheat seeds were surface-sterilized with 1% H_2_O_2_ for 24 h, and then rinsed well with distilled water. The presoaked seeds were placed in culture dishes. After full emergence, the wheat seedlings of uniform size were transplanted to Hoagland’s solution that consisted of the following: 5 mM NH_4_NO_3_, 0.75 mM MgSO_4_, 0.5 mM KH_2_PO_4_, 0.072 mM Fe-EDTA, 0.1 mM Na_2_SiO_3_, 50 μM KCl, 10 μM MnSO_4_, 50 μM H_3_BO_3_, 2 μM ZnSO_4_, 1.5 μM CuSO_4_, and 0.075 μM (NH_4_)_6_Mo_7_O_24_. After 3 days of growth in nutrient solution (pH 6.5), 3 pH levels of nutrient solution were prepared, namely, pH 4.0 (acid stress), pH 6.5 (normal environment as control), and pH 9.0 (alkaline stress); the pH of the solution was adjusted with 1 mol/L HCL and 1 mol/L NaOH. Approximately 10–20 seedlings were measured for each indicator following a completely randomized design. Plants were grown in a growth chamber with a 14-h photoperiod (light intensity 20,000 lux), day/night temperatures of 22°C/19°C, and relative humidity of 64%. The cultivation boxes were randomly moved daily to minimize position effects. The nutrient solution was replaced at the same time every day. Ten-day-old wheat seedlings (second true leaf folded) were selected to monitor net fluxes of H^+^ in roots in response to different rhizosphere pH, and seedlings that were cultured for 15 days were selected for assessment of other indexes.

### Dry matter accumulation

Dry matter accumulation in the roots and shoots at different pH levels was measured by the oven-drying method at 105°C for 30 min and then at 80°C until a constant weight was achieved to determine dry matter. The roots were soaked in 20 mM Na_2_-EDTA for 10 min to remove any metal ions adhering to the root surface before drying.

### Net H^+^ flux at the root surface

To measure H^+^ flux, measuring solution (0.5 mM NH_4_NO_3_, 0.1 mM CaCl_2_, 0.3 mM MES,) with different pH levels were prepared by adjusting the pH with HCl (acidic ingredient) or Tris (alkaline ingredient). This experiment adopted the instantaneous replacement method (quickly replacing testing solutions with different pH during the measurement to change the test pH environment of the material) to measure the absorption kinetics of H^+^ flux at different acid-base levels. Net H^+^ flux was measured using the non-invasive micro-test technology (NMT) (BIO-001A, Younger USA Sci. & Tech. Co., Amherst, MA, USA). The principle behind this method and the instrument used have been previously described [27,28]. Electrodes, holder, and LIX were provided by Xuyue Science & Technology Co. (Beijing, China) (http://www.xuyue.net). The data on H^+^ flux were calculated using MageFlux (Younger USA Corp.; http://voungerusa.com/mageflux).

Microelectrodes were calibrated with different concentrations of H^+^ ion buffers before measurement; electrodes with a slope of -58±2 for measuring H^+^ flux were used in this study. The main radicle, about 3 cm from the root tip, was cut to determine the maximum absorption sites of H^+^ in at least five similar fine roots [29], the maximum ion flux occurred within the region 0 to 15 mm from the root apex. Thus, we selected this region for measurements, balancing 10 min in the measuring solution of pH 6.5 to eliminate local ion imbalances, and then replacing with a new measuring solution for measurement. Next, a blank experiment without plant material was conducted before each formal test to eliminate the influence of the electrode motion agitation of the solution on the ion fluxes [30]. The H^+^ gradient near the root surface (about 5μm above the root surface) was measured by moving the ion-selective microelectrode between two positions separated by a distance of 30 μm in a direction perpendicular to the root axis. To investigate net H^+^ flux at different pH levels, first, ion flux was measured at pH 6.5 for 5 min, and then the test solution was immediately changed to pH 4.0 or pH 9.0 for 10 min. Each treatment was repeated using eight plants, the abnormal data set was removed, and the final calculated ion data were the average of five replicates. The recording rate for the ion flux was one reading per 4 s.

### Root activity

Root activity was assessed using the TTC reduction method. Fifteen seedlings were selected for each treatment, the roots were washed with distilled water, and then the roots were cut to 1-cm-sized cubes and mixed well. Approximately 0.5 g of roots was weighed and placed into a 15-mL centrifuge tube. Each treatment was repeated thrice, 5 mL of phosphoric acid buffer at pH 7.0 and 5 mL of 0.4% TTC were added in succession and kept in the dark at 37°C for 3 h. After incubation, 2 mL of 1 mol/L sulfuric acid were added to stop the reaction. After filtration, 10 mL of methanol were added and incubated at 30°C–40°C for 1-2 h until the root tips were completely white. The OD value was measured at a wavelength of 485 nm using a spectrophotometer.

### Root pH

The roots were cut off and washed with distilled water, and absorbent paper was used to absorb moisture from the root surface. Two grams of fresh roots were ground, and each treatment was repeated thrice. Approximately 4 mL of distilled water were then added and mixed on an oscillator, left to stand at room temperature for 20 min, centrifuged at 12,000 rpm for 15 min, the supernatant was collected and used in measuring pH with a pH meter.

### Data processing

The data shown were analyzed by standard ANOVA assumptions of homogeneity using SPSS 19.0 (Statistical Product and Service Solutions, IBM) and mapping by SigmaPlot 12.0 (Systat Software, Inc., Dundas, CA, USA). Statistical analysis of data was done according to LSD test (p < 0.05). The relationship between H^+^ net flux and root characteristics was assessed by linear correlation analysis.

## Results

### Dry matter accumulation

Under different pH conditions, root dry weight was significantly different between the two cultivars, but the difference in shoot dry weight was not significant (Fig 1). The shoot and root biomass of the two cultivars could be arranged in decreasing order as follows: pH 6.5 > pH 9.0 > pH 4.0. The root dry weight of BN4199 was significantly higher than AK58 under different pH conditions, and shoot dry weight was higher than AK58, but the difference was not significant. The root/shoot ratio of the two varieties increased compared with the control at pH 9.0 and decreased compared with the control at pH 4.0. There was no significant difference in the root/shoot ratio between the two cultivars at pH 6.5, and the root/shoot ratio of BN4199 was significantly greater than AK58 at pH 4.0 and 9.0.

**Fig. 1.**
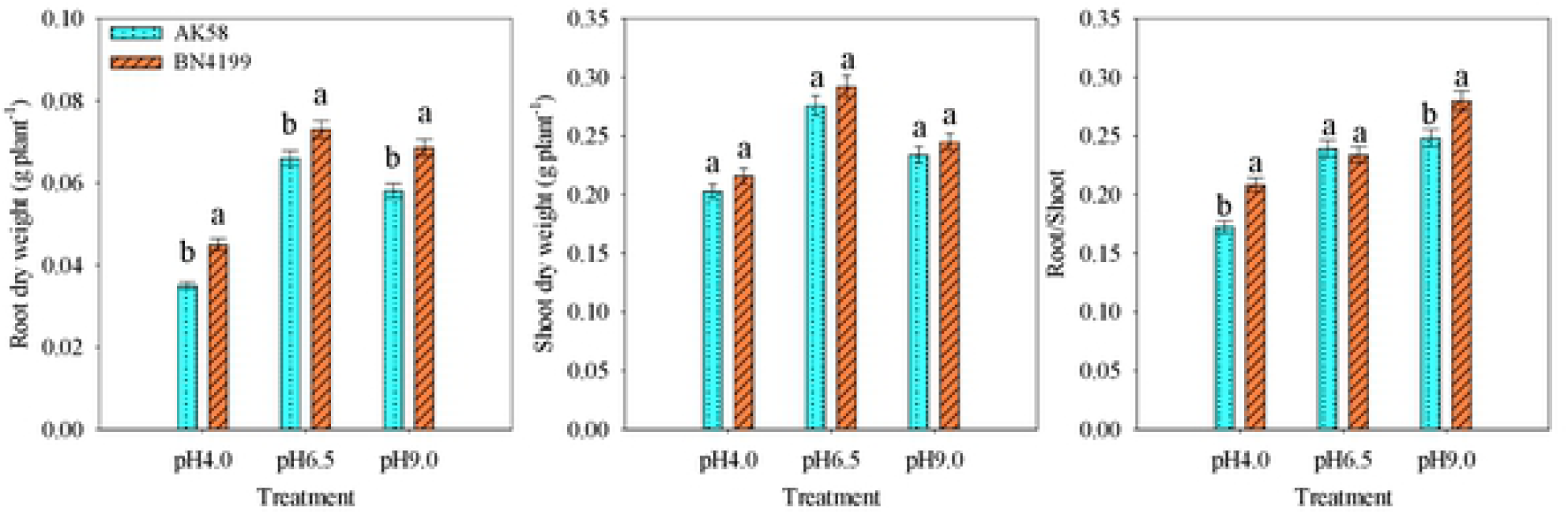
Root and shoot dry weigh of AK58 and BN4199 seedlings afier 15 d of treatment at different pH levels. Results are shown as means ± SD. Vertical bars indicate the standard deviation, and different letters above the error bars indicate significant differences among pH levels between wheat cultivars at P < 0.05 using the LSD test.

### H^+^ net flux

The H^+^ net flux of both cultivars showed efflux at pH 6.5 (Fig 2B). At pH 4.0, the H^+^ net flux of the two varieties significantly decreased compared with the control, whereas H^+^ net flux slowly decreased with time, and the H^+^ net efflux rate approached zero (Fig 2A). At pH 9.0, the instantaneous performance showed a large H^+^ net influx and gradually showed H^+^ net efflux over time (Fig 2C). The change trend of the two wheat varieties at different pH levels was the same. The H^+^ efflux rate of BN4199 was significantly lower than AK58 at pH 6.5 and significantly higher than AK58 at pH 4.0 and 9.0 (Fig 2D).

**Fig. 2.**
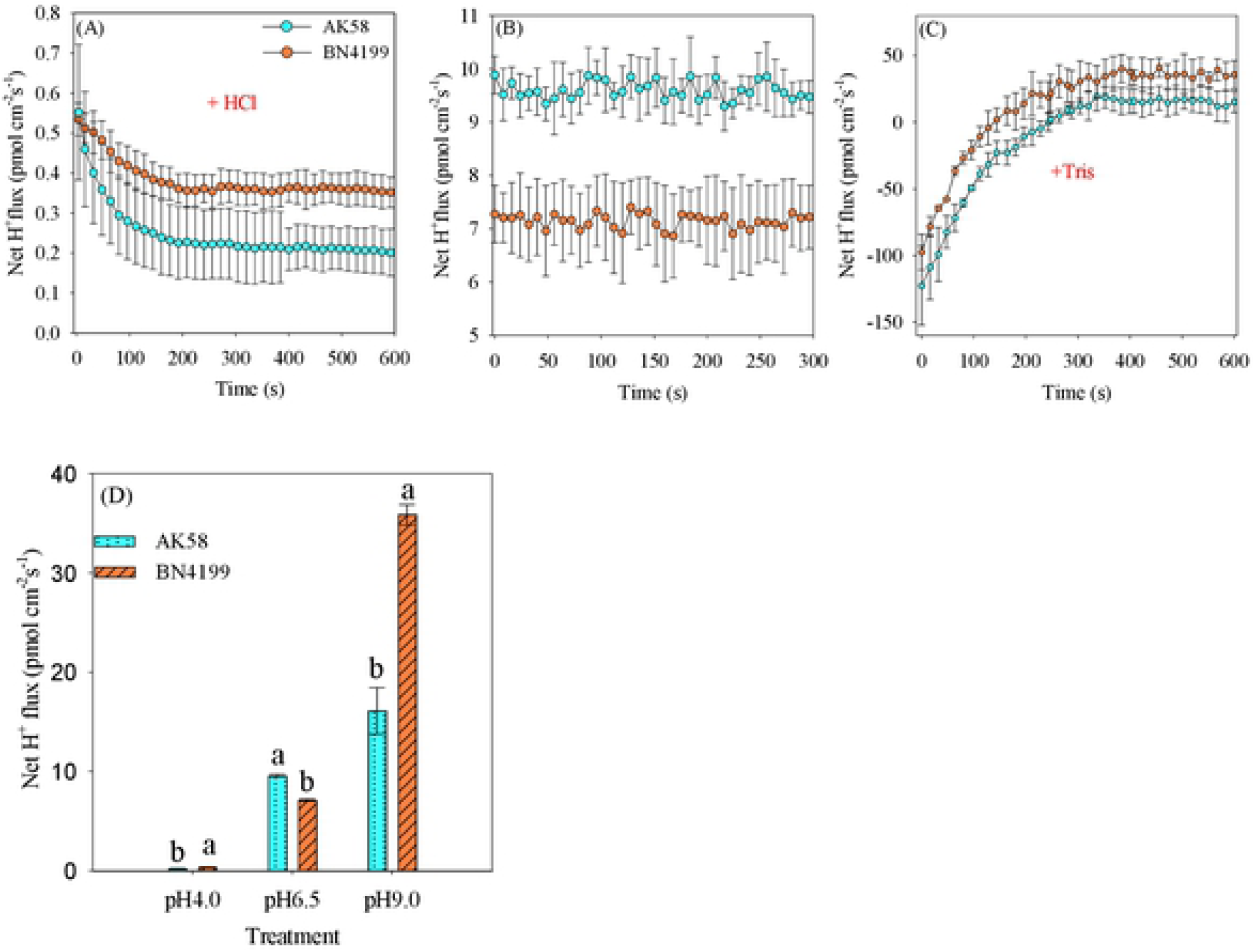
Influence of pH on H ^+^ net flux in wheat root surfaces. Transient H^+^ kinetics of different pH levels in AK58 and BN4I99 roots were measured in root epidermal cells; the dynamic changes in wheat root H^+^ net fluxes (about600 s) are presented after changing acidic test liquid (A) or alkaline test liquid (C). The mean ± SE of H^+^ net fluxes during the measurement period are shown (n = 5). B shows changes in H^+^ net flux after 5 min at pH 6.5. D shows the mean flux of each treatment 1 min before the end of measurement. Vertical bars indicate the standard deviation, and different letters above the error bars indicate significant differences among pH levels between wheat cultivars at P < 0.05 using the LSD test.

### Root activity

The root activity of the two wheat varieties increased with higher pH (Fig 3). There was no significant difference in root activity between the two varieties at pH 6.5 and 9.0, and the root activity of BN4199 was significantly higher than AK58 at pH 4.0.

**Fig. 3.**
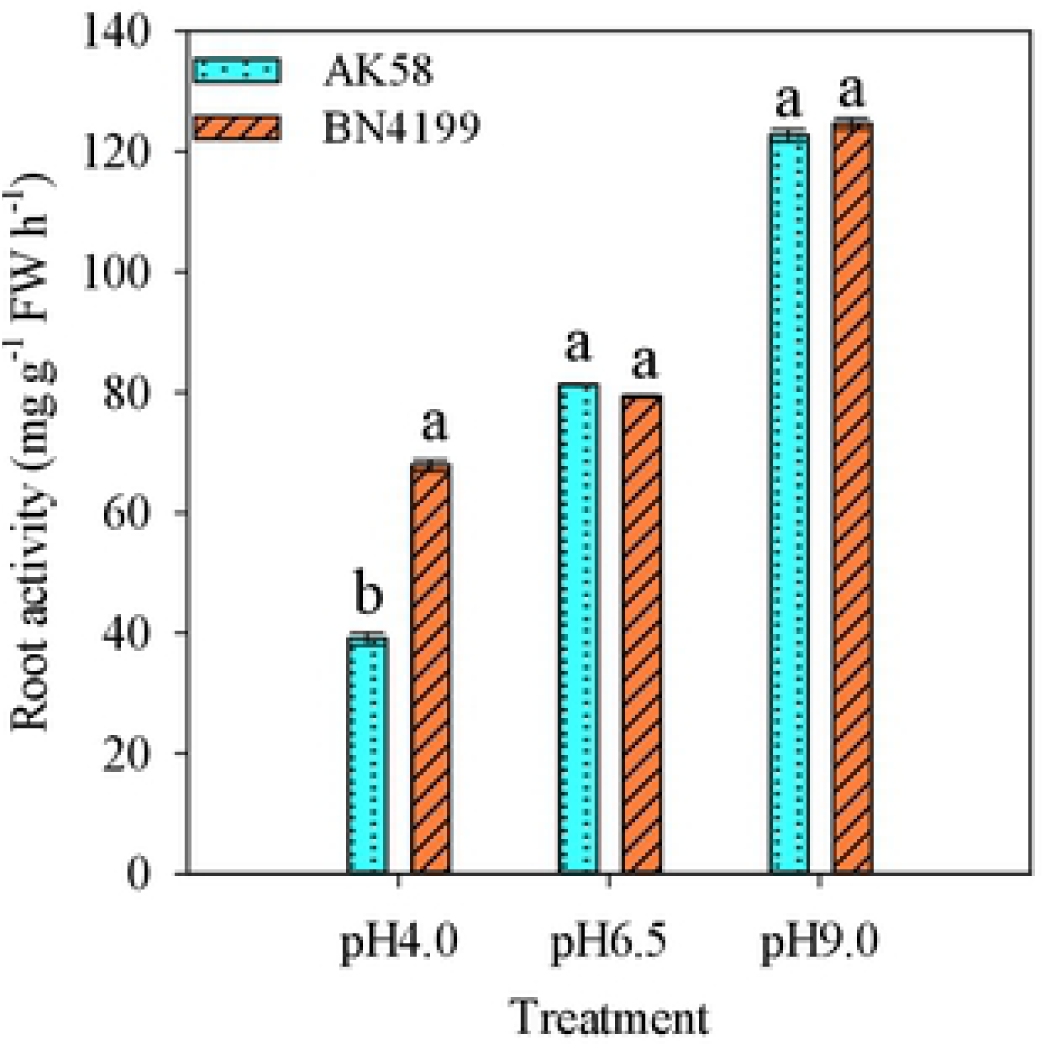
Root activity of AK58 and BN4199 seedlings after 15 d of treatment at different pH levels. Results are shown as means ± SD. Vertical bars indicate the standard deviation, and different lellers above the error bars indicate significant differences among pH values between wheat cultivars at P < 0.05 using the LSD lest.

### Root pH

The root pH of the two varieties was significantly different with various pH levels (Fig 4). The change trend of root pH of the two varieties was similar under different pH conditions and both increased in the following order: pH 4.0 < pH 6.5 < pH 9.0. The root pH of BN4199 was lower than AK58 at pH 6.5 and higher than AK58 at pH 4.0, but the difference was not significant. The root pH of BN4199 was significantly lower than AK58 at pH 9.0. Root pH in BN4199 and AK58 decreased by 1.92% and 2.70% respectively compared to the control at pH 4.0 and increased by 3.99% and 4.15% respectively compared to the control at pH 9.0, indicating that the pH change range of BN4199 was smaller than AK58 under acid and alkali stress.

**Fig. 4.**
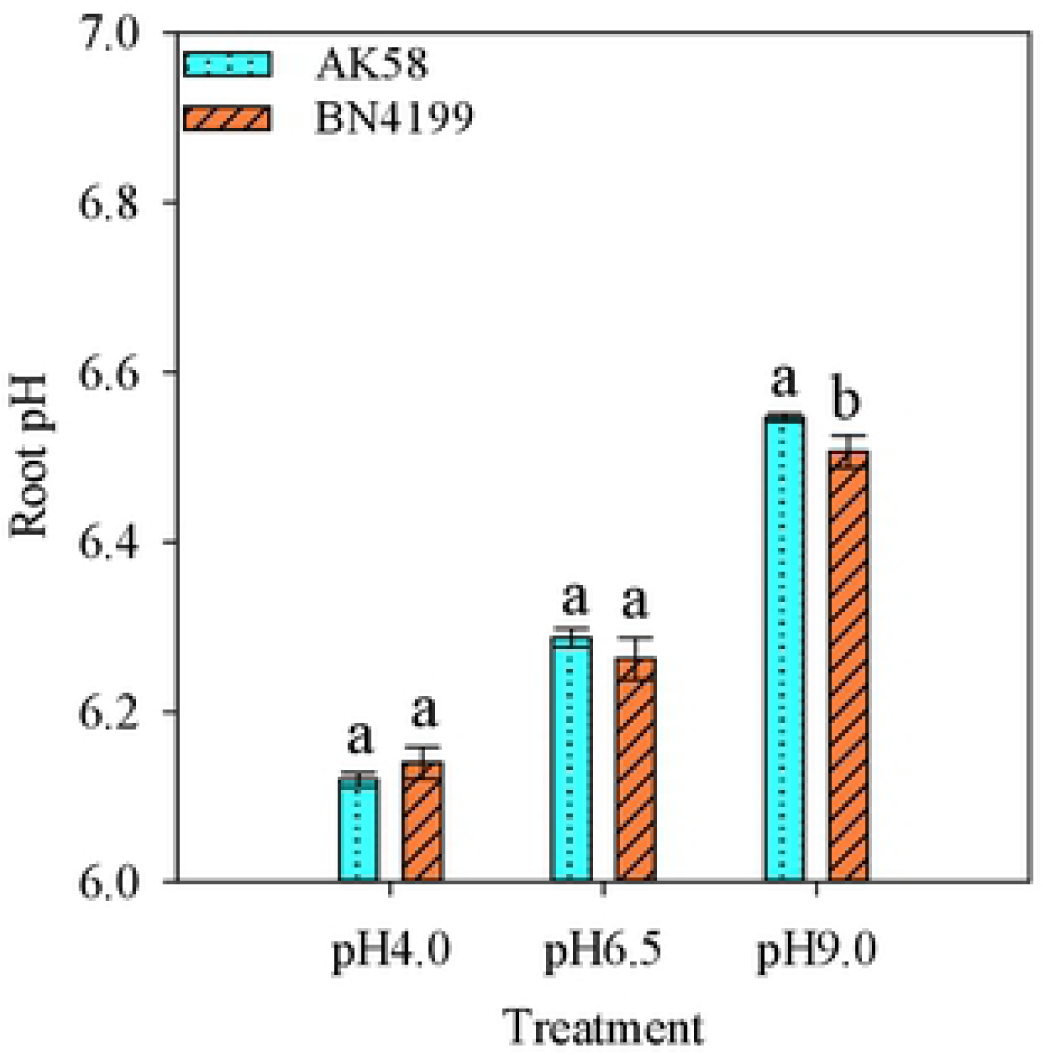
The root pH of AK58 and BN4199 seedlings after 15d of treatment at different pH levels. Results are shown as means± SD. Vertical bars indicate the standard deviation, and different letters above the error bars indicate significant differences among pH levels between wheat cultivars at P < 0.05 using the LSD lest.

### Correlation analysis

Table 1 shows that correlation analysis revealed that shoot and root biomass accumulation was positively but not significantly correlated with changes in H^+^ net flux, root activity, and root pH. The change in H^+^ net flux was significantly positively correlated with root activity and root pH.

**Table 1.** Correlations between root and shoot weight, H^+^ net flux, and root characteristics (H^+^ net flux, root activity, root pH, rhizosphere pH) of winter wheat(*p < 0.05)

| | Shoot weight | Root weight | $H^+$ net flux | Root activity | Root pH |
| --- | --- | --- | --- | --- | --- |
| Root weight | 0.90* | — | 0.60 | 0.58 | 0.52 |
| Shoot weight | — | 0.90* | 0.21 | 0.29 | 0.23 |
| $H^+$ net flux | 0.21 | 0.60 | — | 0.86* | 0.85* |

## Discussion

Rhizosphere pH can affect the absorption and utilization of mineral nutrients by wheat roots [31], thereby affecting the morphology and growth of wheat roots. This study has determined that biomass accumulation in the shoots and roots of wheat was highest at neutral pH, acid-base stress inhibited root and shoot growth, especially under acid stress, and biomass accumulation in wheat roots significantly decreased, while root biomass slightly decreased under alkali stress, and biomass accumulation in shoots was less affected by acid and alkali stress (Fig 1). The accumulation of shoot and root biomass changed the size of the root/shoot ratio of the plant. Under acidic stress, root biomass decreased more than the shoot, and root/shoot ratio decreased. Under alkaline stress, root biomass decreases less than the shoot, thus the root/shoot ratio increased (Fig 1). The root/shoot ratio shows the distribution of assimilation between the aboveground and underground organs. Changes in root/shoot ratio are a common response of plants in response to different stresses [32]. Dry matter accumulation and root/shoot ratio of plants under alkali stress were greater than that under acid stress, suggesting that maintaining a higher root/shoot ratio under acid and alkali stress is beneficial to the accumulation of total plant biomass (Fig 1). Compared with the two varieties, BN4199 has a higher root growth rate and root/shoot ratio under acid alkali stress (Fig 1).

The wheat root system is an active absorbing organ, and the strength of root activity reflects the metabolic ability of the root system, which directly influences the growth of aboveground parts [33]. This study has determined that the root activity of wheat remained high at pH 9.0, whereas at pH 4.0, root respiration was weakened, dehydrogenase activity decreased, thus root activity decreased (Fig 3). H^+^ ion concentrations influence the growth of plant roots and soil microorganisms, whereas the root system can release H^+^ or OH^-^ to compensate for the imbalance in anion and cation absorption at the soil-root interface [10]. The release or absorption of H^+^ by roots changes the rhizosphere pH, thereby influencing nutrient availability [34]. The results of the present showed that the H^+^ efflux rate of wheat roots was significantly higher than that of the control at pH 9.0. H^+^ net efflux acidified the rhizosphere and lowered the pH level of the rhizosphere, which could improve the availability of nutrients in the rhizosphere solution, thereby promoting root growth and increasing the root/shoot ratio (Fig 1 and 2). Oliva et al. [35] suggested that proton efflux under low pH conditions avoided the accumulation of toxic H^+^ in the cytoplasm and supported more nutrient acquisition by H^+^-tolerant species. This study found that the H^+^ efflux rate was significantly lower than that of the control at pH 4.0, root activity decreased, root growth was inhibited, and root/shoot ratio decreased, further indicating that acid stress inhibited the net release of H^+^, and maintaining a high net H^+^ release under acidic condition stabilizes cytoplasmic pH (Fig 1–3). Compared with AK58, BN4199 maintained a higher H^+^ efflux rate and root activity under acid and alkali stress (Fig 2 and 3).

Correlation analysis showed that there was a significant positive correlation between changes in H^+^ net flux and root activity, indicating that root activity adjusted H^+^ net flux changes, and the increase in root activity accelerates cellular redox reactions, increases dehydrogenase activity, and stimulates H^+^ efflux (Table 1). Changes in H^+^ net flux will affect the pH of root cells [36], thereby influencing root pH homeostasis and metabolic pathways [37,38]. Correlation analysis also showed that H^+^ net flux was significantly positively correlated with changes in root pH, indicating that changes in H^+^ net flux regulate root pH, thereby affecting the metabolic activity of root cells (Table 1). Li et al. [39] believed that the efficiency of the intracellular pH regulatory system before and after stress reflects the degree of plant resistance to stress. Compared with the control, the root pH decreased under acidic conditions, while increased under alkaline conditions. Under acidic conditions, the passive influx of H^+^ acidified root pH, the H^+^ efflux rate of BN4199 was higher than that of AK58, which resulted in root pH alkaline (Fig 2 and 4). The H^+^ efflux rate of BN4199 was higher than AK58 under alkaline conditions, consequently rhizosphere acidification reduced root pH, and the acid-base balance inside and outside the cells promoted root growth (Fig 2 and 4). In terms of root pH, this was close to neutral regardless of whether the rhizosphere was acidic or alkaline (Fig 4). This may be a regulatory mechanism for the root system to adapt to the acid-base environment, which needs further investigation.

The effect of rhizosphere pH on wheat varied with variety. Gao et al. [41] considered that the accumulation of biomass reflected the response of plants to stress. Under acid-base stress, biomass accumulation, H^+^ net flux, and root activity of BN4199 were all greater than AK58, and root pH tended to be neutral, which indicated that BN4199 has a strong ability to autonomously adjust root pH under acid-base stress (Fig 1–4). Terletskaya et al. [42] earlier reported that the root/shoot ratio of more tolerant wheat varieties increased with stress, whereas that of sensitive varieties decreased with stress. The root/shoot ratio of BN4199 was significantly higher than AK58 under acid-base conditions, and there was no significant difference between the two varieties under neutral conditions, indicating that BN4199 has stronger acid and alkali resistance than AK58 (Fig 1). Therefore, in wheat ecological breeding, the ability of wheat seedlings to tolerate acid and alkali can be quickly identified by measuring the root/shoot ratio.

## Conclusions

Wheat could adapt to poor acid-base environments by adjusting root H^+^ net flux, and wheat cultivars under different rhizosphere pH values in our study showed variations in acid and alkali tolerance-related traits at the seedling stage. Wheat varieties with strong acid and alkali resistance have high H^+^ efflux rates, strong root activity, and neutral root pH. Root H^+^ efflux under alkali stress acidifies the rhizosphere, which is beneficial to root growth and increases root/shoot ratio. In practice, the root/shoot ratio could be used as index for the rapid determination of acid-base tolerance in wheat at the seedling stage.

## Acknowledgments

The National Key Research and Development Plan (2016YFD0101602) and the Key Scientific and Technological Plan Projects in Henan Province (202102110033) supported this study. We thank LetPub (www.letpub.com) for its linguistic assistance during the preparation of this manuscript.

## Author Contributions

**Data curation: Guangtao Wang**.

**Methodology: Weihua Ding**.

**Project administration: Tiezhu Hu**.

**Supervision: Zhengang Ru**.

**Writing – original draft: Guangtao Wang**.

**Writing – review & editing: Suwei Feng**.

## Compliance with ethical standards

### Conflict of interest

The authors have no conflicts of interest to declare.

## References

1. Bhuyan MHMB, Hasanuzzaman M, Mahmud JA, Hossain MS, Bhuiyan TF, Fujita M. Unraveling Morphophysiological and Biochemical Responses of Triticum aestivum L. to Extreme pH: Coordinated Actions of Antioxidant Defense and Glyoxalase Systems. Plants. 2019;8:24. https://doi.org/10.3390/plants8010024

2. Ghimire R, Machado S, Bista P. Soil pH, Soil Organic Matter, and Crop Yields in Winter Wheat-Summer Fallow Systems. Agronomy Journal. 2017;109:706–717. https://doi.org/10.2134/agronj2016.08.0462

3. Guo JH, Liu XJ, Zhang Y, Shen JL, Han WX, Zhang WF, et al. Significant Acidification in Major Chinese Croplands. Science. 2010;327:1008–1010. https://doi.org/10.1126/science.1182570

4. Zhu QC, Liu XJ, Hao TX, Zeng MF, Shen JB, Zhang FS, et al. Modeling soil acidification in typical Chinese cropping systems. ence of the Total Environment. 2018;613–614:1339-1348. https://doi.org/10.1016/j.scitotenv.2017.06.257

5. Hinsinger P. Bioavailability of soil inorganic P in the rhizosphere as affected by root-induced chemical changes: a review. Plant and Soil. 2001;237:173–195. https://doi.org/10.1023/A:1013351617532

6. Novotny EH, De Freitas Maia CMB, De Melo Carvalho MT, Madari BE. Biochar: Pyrogenic Carbon for Agricultural Use - a Critical Review. Revista Brasileira de Ciência do Solo. 2015;39:321–344. https://doi.org/10.1590/01000683rbcs20140818

7. Kavitha B, Reddy PVL, Kim B, Lee BK, Pandey SK, Kim KH. Benefits and limitations of biochar amendment in agricultural soils: A review. Journal of Environmental Management. 2018;227:146–154. https://doi.org/10.1016/j.jenvman.2018.08.082

8. Kost D, Chen LM, Guo XL, Tian YQ, Ladwig K, Dick WA. Effects of flue gas desulfurization and mined gypsums on soil properties and on hay and corn growth in eastern Ohio. J Environ Qual. 2014;43:312–321. https://doi.org/10.2134/jeq2012.0157

9. Feng SW, Gu SB, Zhang HB, Wang D. Root vertical distribution is important to improve water use efficiency and grain yield of wheat. Field Crops Research. 2017;214:131–141. https://doi.org/10.1016/j.fcr.2017.08.007

10. Hinsinger P, Plassard C, Tang CX, Jaillard B. Origins of root-mediated pH changes in the rhizosphere and their responses to environmental constraints: A review.Plant and Soil. 2003;248:43–59. https://doi.org/10.1023/A:1022371130939

11. Cu STT, Hutson J, Schuller KA. Mixed culture of wheat (Triticum aestivum L.) with white lupin (Lupinus albus L.) improves the growth and phosphorus nutrition of the wheat. Plant and Soil. 2005;272:143–151. https://doi.org/10.1007/s11104-004-4336-8

12. Bravin MN, Tentscher P, Rose I, Hinsinger P. Rhizosphere pH Gradient Controls Copper Availability in a Strongly Acidic Soil. Environmental ence & Technology. 2009;43:5686–5691. https://doi.org/10.1021/es900055k

13. Hinsinger P, Bengough AG, Vetterlein D, Young IM. Rhizosphere: biophysics, biogeochemistry and ecological relevance. Plant and Soil. 2009;321:117–152. https://doi.org/10.1007/s11104-008-9885-9

14. Hinsinger P. Plant-induced changes of soil processes and properties. Soil Conditions and Plant Growth. Blackwell Publishing Ltd, Oxford. 2013 pp:323–365. https://doi.org/10.1002/9781118337295.ch10

15. Shavrukov Y, Hirai Y. Good and bad protons: genetic aspects of acidity stress responses in plants. Journal of Experimental Botany. 2015;67:15–30. https://doi.org/10.1093/jxb/erv437

16. Xue ZY, Zhi DY, Xue GP, Zhang H, Zhao YX, Xia GM. Enhanced salt tolerance of transgenic wheat (Tritivum aestivum L.) expressing a vacuolar Na+/H+ antiporter gene with improved grain yields in saline soils in the field and a reduced level of leaf Na+. Plant science. 2004;167:849–859. https://doi.org/10.1016/j.plantsci.2004.05.034

17. Bouthour D, Kalai T, Chaffei HC, Gouia H, Corpas FJ. Differential response of NADP-dehydrogenases and carbon metabolism in leaves and roots of two durum wheat (Triticum durum Desf.) cultivars (Karim and Azizi) with different sensitivities to salt stress. Journal of Plant Physiology. 2015;179:56–63. https://doi.org/10.1016/j.jplph.2015.02.009

18. Garcia-Oliveira AL, Martins-Lopes P, Tolrà R, Poschenrieder C, Guedes-Pinto H, Benito C. Differential Physiological Responses of Portuguese Bread Wheat (Triticum aestivum L.) Genotypes under Aluminium Stress. Diversity. 2016;8:26. https://doi.org/10.3390/d8040026

19. Pena LB, Méndez AAE, Matayoshi CL, Zawoznik MS. Early response of wheat seminal roots growing under copper excess. Plant Physiol Biochem. 2015;87:115–123. https://doi.org/10.1016/j.plaphy.2014.12.021

20. Rasafi TE, Nouri M, Bouda S, Haddioui A. The Effect of Cd, Zn and Fe on Seed Germination and Early Seedling Growth of Wheat and Bean. Ekológia. 2016;35:213–223. https://doi.org/10.1515/eko-2016-0017

21. Sheng HJ, Zeng J, Liu Y, Wang XL, Wang Y, Kang HY, et al. Sulfur Mediated Alleviation of Mn Toxicity in Polish Wheat Relates to Regulating Mn Allocation and Improving Antioxidant System. Frontiers in Plant ence. 2016;7:1382. https://doi.org/10.3389/fpls.2016.01382

22. Wang HO, Zhong GG, Shi GQ, Pan FT, et al. Toxicity of Cu, Pb, and Zn on Seed Germination and Young Seedlings of Wheat (Triticum aestivum L.). Computer & Computing Technologies in Agriculture Iv-ifip Tc 12 Conference. DBLP. 2011. https://doi.org/10.1007/978-3-642-18354-6_29

23. Mathur S, Kalaji HM, Jajoo A. Investigation of deleterious effects of chromium phytotoxicity and photosynthesis in wheat plant. Photosynthetica. 2016;54:185–192. https://doi.org/10.1007/s11099-016-0198-6

24. Carillo P, Annunziata MG, Pontecorvo G, Fuggi A, Woodrow P. Salinity Stress and Salt Tolerance. Abiotic Stress in Plants-Mechanisms and Adaptations. Intech Open, London. 2011;pp:21-38. https://doi.org/10.5772/22331

25. Borzouei A, Kafi M, Akbari-Ghogdi E, Mousavi-Shalmani MA. Long term salinity stress in relation to lipid peroxidation, super oxide dismutase activity and proline content of saltsensitive and salt-tolerant wheat cultivars. Chilean journal of agricultural research. 2012;72:476–482. https://doi.org/10.4067/S0718-58392012000400003

26. Tang C, Nuruzzaman M, Rengel Z. Screening wheat genotypes for tolerance of soil acidity. Australian Journal of Agricultural Research. 2003;54:445–452. https://doi.org/10.1071/AR02116

27. Wherrett T, Ryan PR, Delhaize E, Shabala S. Effect of aluminium on membrane potential and ion fluxes at the apices of wheat roots. Functional Plant Biology. 2005;32:199–208. https://doi.org/10.1071/FP04210

28. Shabala S. Non-Invasive Microelectrode Ion Flux Measurements In Plant Stress Physiology. In: Volkov A.G. (eds) Plant Electrophysiology. Springer, Berlin. 2006;pp:35-71. https://doi.org/10.1007/978-3-540-37843-3_3

29. Wu HH, Zhu M, Shabala L, Zhou MX, Shabala S. K+ retention in leaf mesophyll, an overlooked component of salinity tolerance mechanism: A case study for barley. Journal of Integrative Plant Biology. 2015;57:171–185. https://doi.org/0.1111/jipb.12238

30. Zhang CX, Meng S, Li YM, Zhao Z. Net NH4+ and NO3-fluxes, and expression of NH4+ and NO3-transporter genes in roots of Populus simonii after acclimation to moderate salinity. Trees. 2014;28(6):1813–1821. https://doi.org/10.1007/s00468-014-1088-9

31. Mak M, Babla M, Xu SC, O’Carrigan A, Liu XH, Gong YM, et al. Leaf mesophyll K+, H+ and Ca2+ fluxes are involved in drought-induced decrease in photosynthesis and stomatal closure in soybean. Environ. Exp Bot. 2014;98:1–12. https://doi.org/10.1016/j.envexpbot.2013.10.003

32. Baquy AMA, Li JY, Xu CY, Mehmood K, Xu RK. Determination of critical pH and Al concentration of acidic Ultisols for wheat and canola crops. Solid Earth. 2017;8:149–159. https://doi.org/10.5194/se-8-149-2017

33. Agathokleous E, Belz RG, Kitao M, Koike T, Calabrese EJ. Does the root to shoot ratio show a hormetic response to stress? An ecological and environmental perspective. Journal of Forestry Research. 2019;30:1569–1580. https://doi.org/10.1007/s11676-018-0863-7

34. Zhou Y, Yang XW, Zhou SM, Wang YJ, Yang R, Xu DF, et al. Activities of Key Enzymes in Root NADP-Dehydrogenase System and Their Relationships with Root Vigor and Grain Yield Formation in Wheat. Scientia Agricultura Sinica. 2018;51:2060–2071. https://doi.org/10.3864/j.issn.0578-1752.2018.11.004

35. Custos JM, Moyne C, Sterckeman T. How root nutrient uptake affects rhizosphere pH: A modelling study. Geoderma. 2020;369:114314. https://doi.org/10.1016/j.geoderma.2020.114314

36. Oliva SR, Mingorance MD, Sanhueza D, Fry SC, Leidi EO. Active proton efflux, nutrient retention and boron-bridging of pectin are related to greater tolerance of proton toxicity in the roots of two Erica species. Plant Physiology and Biochemistry. 2018;126:142–151. https://doi.org/10.1016/j.plaphy.2018.02.029

37. Zhang YK, Zhu DF, Zhang YP, Chen HZ, Xiang J, Lin XQ. Low pH-induced changes of antioxidant enzyme and ATPase activities in the roots of rice (Oryza sativa L.) seedlings. PloS One. 2015;10:e0116971. https://doi.org/10.1371/journal.pone.0116971

38. Neumann G, Römheld V. Rhizosphere Chemistry in Relation to Plant Nutrition. Marschner’s Mineral Nutrition of Higher Plants (Third Edition), Academic Press, San Diego. 2012;pp:347–368. https://doi.org/10.1016/B978-0-12-384905-2.00014-5

39. Inoue SI, Takahashi K, Okumura-Noda H, Kinoshita T. Auxin Influx Carrier AUX1 Confers Acid Resistance for Arabidopsis Root Elongation Through the Regulation of Plasma Membrane H+-ATPase. Plant Cell Physiol. 2016;57:2194–2201. https://doi.org/10.1093/pcp/pcw136

40. Li XB, Pan JW, Ding GH, Li Z, Wang QG, Liu W. Determination of pH Value in Root Cells of Foxtail Millet under Salt Stress. Shandong Agricultural Sciences. 2019;51:40–44. https://doi.org/CNKI:SUN:AGRI.0.2019-02-008

41. Gao ZW, Han JY, Mu CS, Lin JX, Li XY, Lin LD, et al. Effects of Saline and Alkaline Stresses on Growth and Physiological Changes in Oat (Avena sativa L.) Seedlings.Notulae Botanicae Horti Agrobotanici Cluj-Napoca. 2014;42:357–362. https://doi.org/10.15835/nbha4229441

42. Terletskaya N, Rysbekova A, Iskakova A, Khailenko N, Polimbetova F. Saline stress response of plantlets of common wheat (Triticum aestivum) and its wild congeners. J Agr Sci Tech B. 2011;1:198–204.

43. Chaignon V, Quesnoit M, Hinsinger P. Copper availability and bioavailability are controlled by rhizosphere pH in rape grown in an acidic Cu-contaminated soil. Environ Pollut. 2009;157:3363–3369.

